# The use of Benzonase to produce ribosome footprints simplifies translational levels quantification by Ribo-seq

**DOI:** 10.1101/2025.03.07.642103

**Authors:** Guillermo Eastman, George S. Bloom, José R. Sotelo-Silveira

**Affiliations:** Department of Biology, University of Virginia, Charlottesville, VA, 22904, USA; Department of Neuroscience, University of Virginia, Charlottesville, VA, 22903, USA; Department of Cell Biology, University of Virginia, Charlottesville, VA, 22903, USA; Departamento de Genómica, Instituto de Investigaciones Biológicas Clemente Estable, Ministerio de Educación y Cultura, Montevideo, 11600, Uruguay; Sección Biología Celular, Facultad de Ciencias, Universidad de la República, Montevideo, 11400, Uruguay

## Abstract

Gene expression quantification through genomics methods is crucial for understanding diverse biological contexts. Among these methods, ribosome profiling (Ribo-seq) stands out as a valuable tool for uncovering post-transcriptional gene expression regulation by providing a comprehensive view of the translatome. While current protocols are time-intensive with limited variations, we introduced the use of the Benzonase enzyme to generate ribosome footprints from a polysome-enriched fraction that exhibit expected characteristics in size, transcriptome mapping, and periodicity. Comparing translatome from Benzonase- and RNAse I-derived footprints reveals minimal differences underscoring Benzonase’s potential to streamline the protocol, reducing time, cost and bias. We further demonstrate Ribo-seq application in primary neuronal cultures, using Benzonase to digest the post-mitochondrial supernatant, thereby bypassing the labor-intensive ribosome/polysome purification step. The introduction of such protocol variations for Ribo-seq, especially for challenging low-input samples, offers a significant advancement of this resource for the community.

**GRAPHICAL ABSTRACT:** 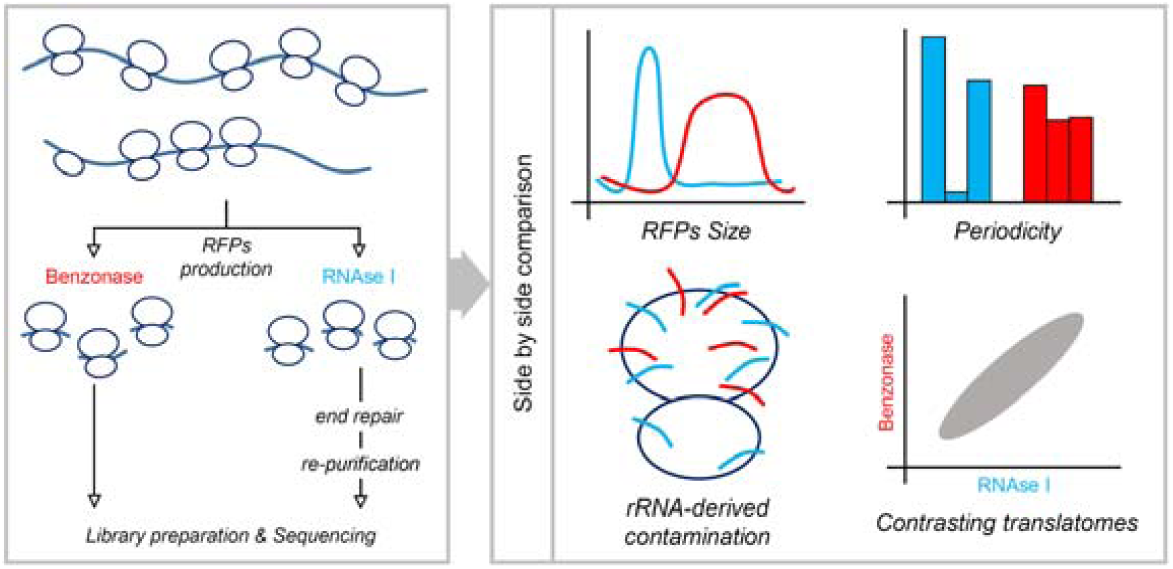

## INTRODUCTION

The comprehensive genome-wide study of gene expression serves as an indispensable instrument used to understand how biological systems respond to different stimuli, pharmacological agents, or environmental variables. Within this scientific purview, numerous experimental approaches and techniques have been developed to unravel the regulatory mechanisms that control each step in the gene expression process. Centrally to this discussion the ribosome profiling protocol, termed Ribo-seq (1), facilitates an in-depth examination of ribosomal occupancy on mRNA molecules on a genome-wide scale, achieving a remarkable subcodon resolution. Importantly, by quantifying the ribosomal footprints on a specific mRNA, it becomes feasible to derive an estimate of the translation efficiency of said mRNA (see the following reviews for more details (2, 3)). From a broader genomic perspective, such analysis delineates the “translatome”, a new layer of quantitative information encompassing several gene expression regulation mechanisms, including post-transcriptional regulation which can be lost quantifying just the transcriptome.

The Ribo-seq protocol, while intricate and time-intensive, can be broadly summarized in three primary steps: i) isolation of a translational cell sample, which includes cell lysate and polysome collection; ii) RNAse digestion followed by the recovery of ribosome footprints; and iii) deep sequencing and in silico analysis. Briefly, the foundational protocol (4) begins with the cell lysis, proceeds to the RNAse digestion assay, and then to the isolation of ribosomes via ultracentrifugation. Subsequently, ribosome footprints, ranging between 26-34 nucleotides in size, are recovered from a gel electrophoresis. Here, we introduce adjustments to this protocol to enhance the translational readout. Basically, after cell lysis, we isolate the polysomes by ultracentrifugation (5) before performing the RNAse digestion step. This approach aims to increase confidence in the translational status of the ribosomes generating the footprints, mitigating potential scenarios where inactive ribosomes or other ribonucleoparticles might produce ambiguous footprints. Other methodological alternatives have recently been described to assure proper translatome quantification, particularly in low-input samples. For example, the use of a template switch retrotranscriptase to produce the DNA libraries avoids ligation steps and ensures results from ultra low-input samples (6). On the other hand, other groups have shown that ribosome footprints can be obtained directly from the cell lysate bypassing the ribosome isolation step by ultracentrifugation (7, 8).

The controlled RNase digestion step is pivotal in the Ribo-seq protocol, serving as a primary factor in ribosome footprint production (9–11). In this context, the RNAse I enzyme has been the most widely utilized, as the one described in the original protocol (4). However, a few other enzymes have been described and used in the literature, such as micrococcal nuclease (12), and RNAse A, S7 and T1 (9). In this work, we introduce the use of the Benzonase nuclease. While there are variations in the features between the RNAse I and Benzonase enzymes, for example, the enzymatic activity over single or double RNA strands or DNA, one is critical for the practical application of the Ribo-seq protocol, such as the phosphorylation status of the 5’ and 3’ end in their degraded fragments. Specifically, RNAse I generates fragments with a phosphate group at the 3’ end, whereas Benzonase incorporates a phosphate group at the 5’ end. We capitalize this slight difference in the ligation step after the ribosome footprints isolation step: Benzonase-derived footprints are ready to be ligated, while RNAse I-derived fragments need to be simultaneously dephosphorylated at the 3’ end and phosphorylated at the 5’ end. This adds at least one enzymatic reaction and one RNA precipitation and recovery step to the protocol. Such additions not only have the potential to introduce biases but may also diminish the final RNA yield. Moreover, the 5’-end phosphorylation needed for the RNAse I-derived footprints, will not distinguish between genuine ribosome footprints and degraded RNA fragments, both dephosphorylated at the 3’-end. Therefore, noise can be introduced in the sequencing data. Conversely, by employing Benzonase we mitigate the inclusion of such degraded fragments.

In this study, we sought to compare ribosome footprints generated using either Benzonase or RNAse I and optimize the Benzonase protocol to be used in low-input type situations, such as primary cell cultures. We contrasted global features of the ribosome footprints, including their length, mapping distribution over mRNAs and periodicity. Additionally, we examined rRNA fragments produced by both digestion methods and compared the translatomes defined by each enzyme. We investigated individual genes that were overrepresented in one translatome or associated with the use of one enzyme and found a particular bias related to coding sequence (CDS) length. Although we observed some expected differences in the ribosome footprint characteristics, such as broader length distribution for Benzonase-derived footprints and a distinct periodicity pattern in RNAse I-derived footprints, the translatomes defined by both enzymes were highly comparable.

Finally, following previous modifications reported in the literature, we optimized the Ribo-seq protocol to be applicable in low-input cellular models, such as embryonic neuronal cultures. We eliminated the ultracentrifugation step to recover either ribosomes or polysomes and performed the RNAse digestion assay with Benzonase directly on a post-mitochondrial supernatant. To demonstrate the efficacy of this modified protocol, we extracted and sequenced ribosome footprints from two mouse embryonic neuronal cultures (4 to 6 million cells), revealing the anticipated primary characteristics of both the ribosome footprints and translatomes.

Considering the extensive use of Ribo-seq, we think that new methodological variations are necessary to adjust the technique to unique biological models and specific research questions. The use of Benzonase to produce ribosome footprints would be a valuable resource for the community exploring translation regulation, but in particular for new research groups without advanced expertise background. Also, the continuous development of new protocols, such as single-cell approaches, would require a complete understanding of the features of the available enzymes to produce ribosome footprints, as the one described here for the Benzonase.

## MATERIAL AND METHODS

### Cell line and neuronal cultures

The cell line HEK293 was grown in DMEM (Gibco, Cat# 10313021) with 10% of calf serum (Cytiva, Cat# SH30087.03) and antibiotics (Sigma-Aldrich, Cat# A5955), at 37°C and 5% of CO_2_ in a humid atmosphere. Primary neuron cultures were obtained from E17-18 brain cortices dissected from wild-type (C57/Bl6) or tau knock-out mice (13), as previously described (14). Neuron cultures were grown in poly-D-lysine (Sigma, Cat# P0899) coated surfaces for 10-11 days in vitro in Neurobasal medium (Gibco, Cat# 21103049) supplemented with B-27 (Gibco, Cat# 17504044) and in the presence of 1.5 g/L of glucose (Sigma, Cat# 16301), 2 mM of GlutaMAX (Gibco, Cat# 35050061) and 10 µg/mL of gentamicin (Gibco, Cat# 15710064).

### Ribo-seq protocol

Cultures were pretreated for 1 hour at 37°C with 100 µg/mL of cycloheximide (CHX; Sigma-Aldrich, Cat# 01810) to stop translation, then kept on ice and washed three times with cold PBS and 100 100 µg/mL of CHX. Cells are detached mechanically using a scraper, centrifuged for 10 minutes at 2000 RPM and 4°C, resuspended and lysed in the presence of 5 mM Tris pH 7.5; 2.5 mM MgCl_2_; 1.5 mM KCl; 0.5% Triton X-100; 0.5% sodium deoxycholate; 2 mM DTT and 100 µg/mL CHX. Post-mitochondrial supernatant (PMS) was obtained by centrifugation at full speed (17, 000 g) for 2 minutes at 4°C. For human-derived cell lines, PMS was loaded in 12-33.5% sucrose cushion in buffer containing 20 mM HEPES pH 7.5; 5 mM MgCl_2_; 100 mM KCl y 100 µg/mL CHX and ultracentrifuged using a Beckman SW41Ti rotor for 2 hours at 36, 000 RPM and 4°C. The obtained polysomal pellet was resuspended in the same sucrose buffer and used for subsequent RNAse protection assays. For neuronal cultures, RNAse treatment was directly performed over PMS.

Polysome fractions were digested either with 200 U of Benzonase nuclease (Sigma-Aldrich, Cat# E8263) for 10 minutes at room temperature (RT) or with 300 U of RNAse I (Invitrogen, Cat# AM2294) for 45 minutes at RT with gentle mixing. In both cases, digestion was interrupted by adding 3 volumes of mirVana Lysis Buffer and then continued with RNA isolation using the mirVana kit (Invitrogen, Cat# AM1560). The eluted RNA was precipitated with 80% ethanol and 3M sodium acetate pH 5.2 to maximize small RNA recovery and then dissolved in RNAse-free water. Digested RNA was denatured at 80°C for 90 seconds and then separated using 15% PAGE in TBE buffer (89 mM Tris-borate pH 8.3 and 2 mM EDTA) and in the presence of 7 M Urea (Invitrogen, Cat# EC68852BOX) for 65 minutes at 200 V. Gels were revealed using GelRed (Biotium, Cat# 41003) and visualized in a dark room under UV light exposure. Ribosome footprints bands were recognized using two synthetic ssRNA of 26 and 34 nucleotides (nt) in length (Eurofins) and an ultra-low range DNA ladder (Invitrogen, Cat# 10597012). Bands were sliced out and RNA was extracted by overnight mixing with 0.3 M sodium acetate pH 5.5; 1 mM EDTA and 0.25% (wt/vol) SDS. RNA was precipitated overnight at -20°C using isopropanol and resuspended in RNAse-free water for quality control before shipment to sequencing. In the case of RNAse I-derived ribosome footprints, before quality control, RNA was simultaneously phosphorylated at 5’-ends and dephosphorylated at 3’-ends by the action of 10 U of T4 Polynucleotide Kinase (New England Biolabs, Cat# M0201) in the presence of 1 mM of ATP for 30 minutes at 37°C. Ribosome footprints were then precipitated with 0.3 M sodium acetate and 80% ethanol, concentrated and resuspended in RNAse-free water.

Quality control of ribosome footprints was performed by capillary electrophoresis using small RNA chips and the 2100 Agilent Bioanalyzer instrument. BGI Tech Solutions (Hong Kong) performed sequencing using a small RNA library protocol and single-end (50 bp) reads, yielding an average of 120 million reads. Raw sequencing data are available at the NCBI Sequence Read Archive (SRA) under BioProject IDs PRJNA1220030 and PRJNA1220074.

### Data analysis

Briefly, sequence quality was analyzed using FastQC (15) and trimmed, if necessary, using sickle (16). Mapping was performed using bowtie2 (17) and default parameters against the latest version available of mouse and human reference genomes (Mus musculus, mm10 version - GRCm38; and Homo sapiens, hg38 version - GRCh38, respectively) downloaded from the NCBI RefSeq ftp website. Read counts tables were obtained using featureCounts (18).

For the identification of genes specifically detected or abundant with one enzyme, we defined the two following criteria: i) select those genes that change more than a quartile in the expression ranking, and ii) select the top100 genes with the highest difference in gene expression (CPKM) between the enzyme-derived translatomes. Downstream analyses were performed using bash, python and command-line tools as samtools (19) and bedtools (20). Several R packages were used for graph construction and data visualization.

Ribosome structure was obtained from (21) and visualized using the Protein Data Bank (PDB ID: 6QZP) (22).

## RESULTS

### Benzonase and RNAse I produce comparable ribosome footprints obtained from a polysome-enriched fraction

To compare the ribosome footprints produced with either Benzonase or RNAse I enzymes, we isolated a polysome-enriched fraction from HEK293 cell cultures and digested it with both enzymes in parallel. As anticipated, the polysome digestion profiles differed, as indicated by variations in band patterns and intensities. These differences were detected using 15% polyacrylamide gel electrophoresis (PAGE) under the presence of 7 M Urea and further validated through capillary electrophoresis (Fig. 1). Notably, the ribosomal footprint region, approximately 30 nucleotides in length, exhibited subtle distinctions between the two enzymes. High-resolution analysis using Bioanalyzer small RNA chips revealed that Benzonase and RNAse I-generated footprints differ slightly in size and quantity (Fig. 1B).

**Figure 1:**
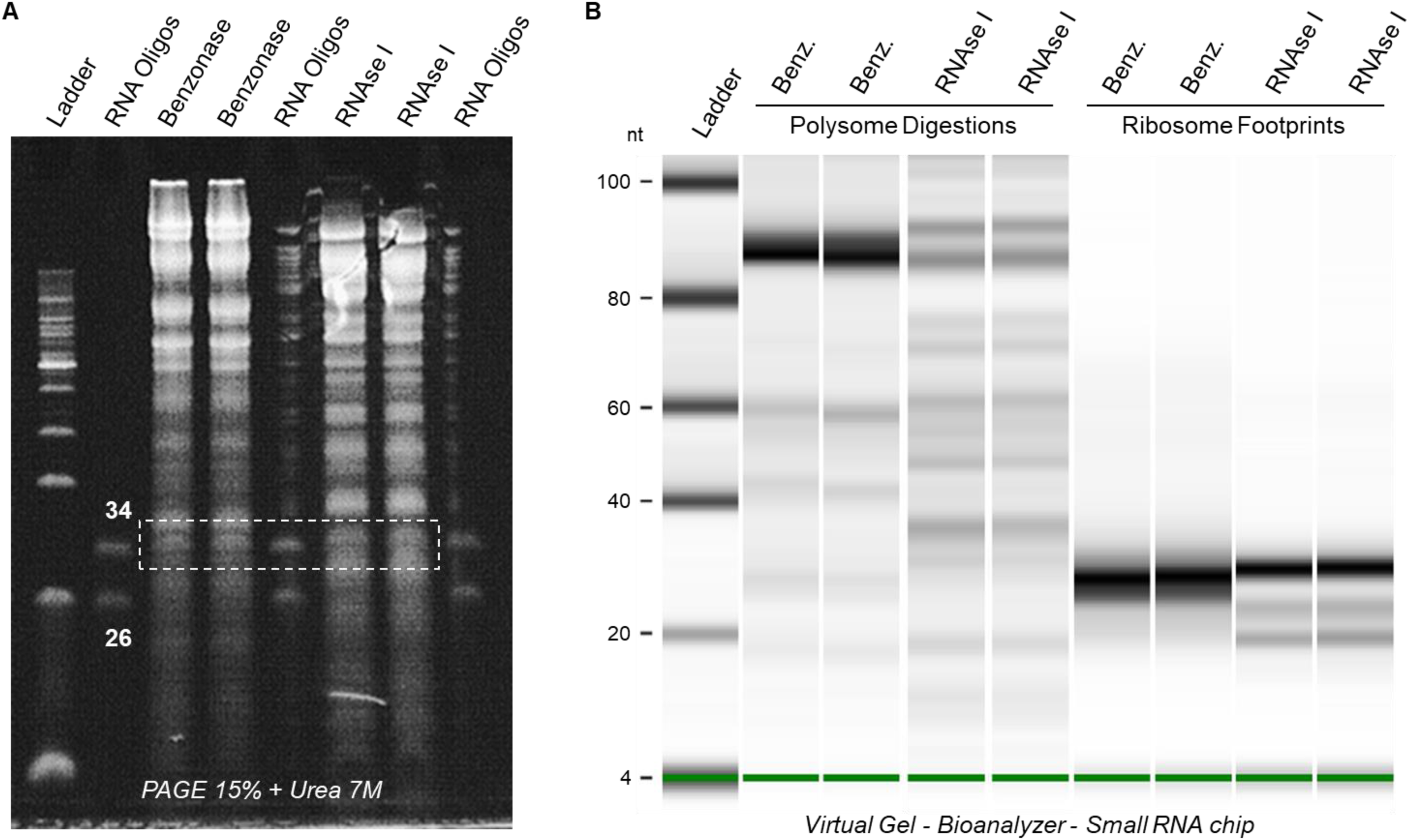
Ribosome footprints production and isolation using Benzonase and RNAse. **I.** A polysome-enriched fraction obtained by ultracentrifugation in sucrose cushions was used to produce ribosome footprints by a ribosome protective assay using either Benzonase or RNAse I enzymes. (A) Digested polysome samples were separated in 15% PAGE in the presence of 7 M Urea. Two ssRNA oligos of 26 and 34 nt in length were used to identify and slice out the gel region that corresponds to the ribosome footprints, indicated by a dotted white box. (B) Digested polysome samples were also analyzed using capillary electrophoresis (Agilent - Bioanalyzer small RNA chips) to visualize RNA population in terms of size and quantity. A virtual reconstruction of the gel is shown for both polysome digestions and the isolated ribosome footprint samples obtained with Benzonase or RNAse I, as indicated.

### Main ribosome footprint features are present in Benzonase-derived footprints

After purifying ribosome footprints we proceed with deep high-throughput sequencing, obtaining 120 million single-end reads per sample on average. Approximately 85% of these reads were derived from rRNA fragments, as detailed in Table S1. Nonetheless, we successfully mapped over 10 million reads to the reference genome, with an average of 85% of these reads corresponding to mRNA regions, as indicated in Table S1.

In our initial comparative analysis of ribosome footprints derived from Benzonase and RNAse I, we examined multiple characteristics of ribosome footprints. These characteristics include the distribution of reads across various regions of mRNA, the length of the footprints, and the periodicity of the reads, as illustrated in Figures 2 and 3.

**Figure 2:**
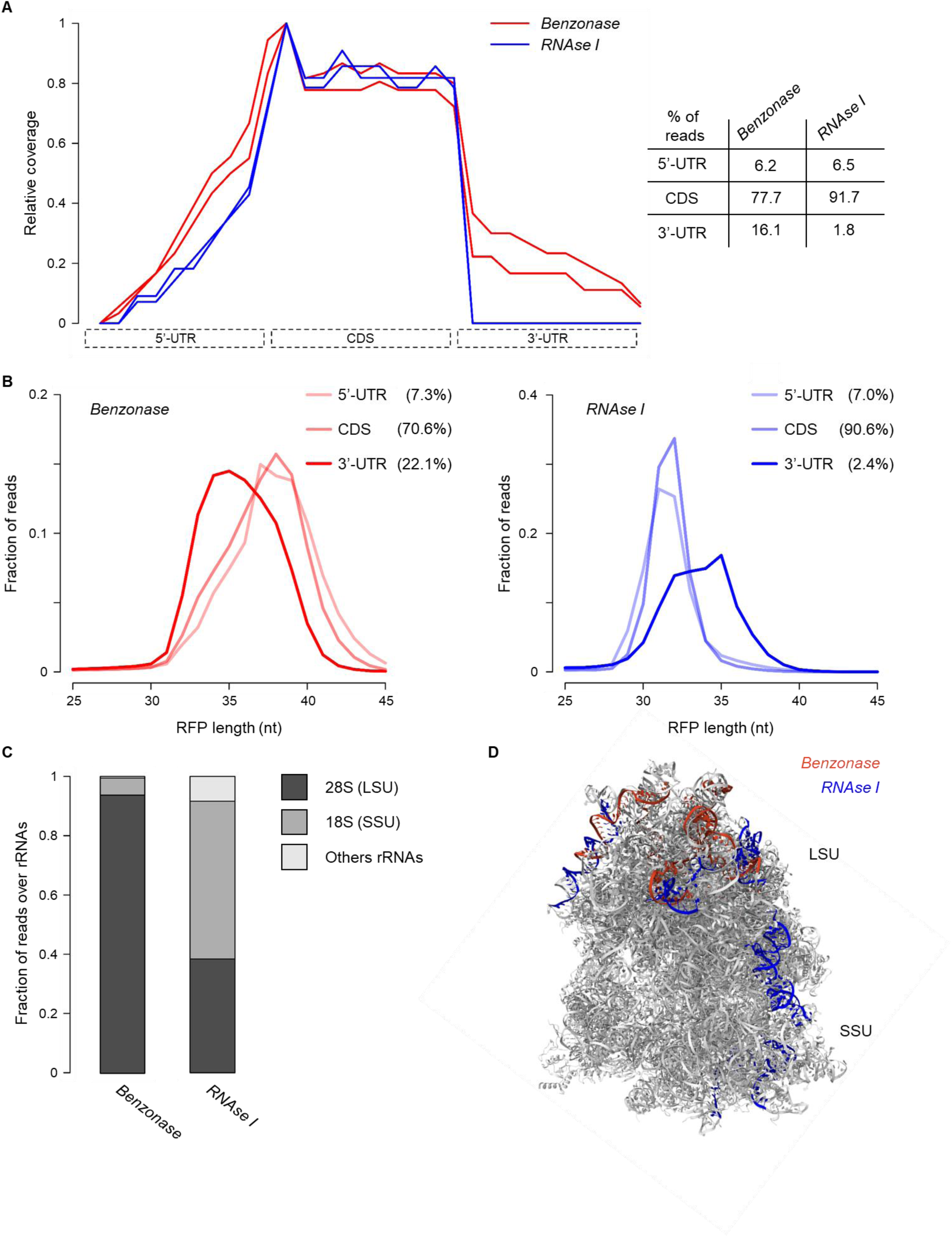
Ribosome footprints mapping distribution among mRNA and rRNA. (A) Global representation of the ribosome footprints coverage over mRNA main regions with Benzonase- and RNAse I-derived footprints. Both enzymes produce footprints mainly associated with coding regions, but a higher proportion of reads mapping at the 3’-UTR is observed for Benzonase-derived footprints compared to RNAse I-derived. (B) Ribosome footprint length distribution is shown for each mapping region (5’-UTR, CDS and 3’-UTR) for Benzonase (left panel, red) and RNAse I (right panel, blue). For each enzyme, the size of the footprints mapping over 5’-UTR and CDS is similar, while footprints mapping over 3’-UTR have a different size distribution. (C) Reads mapping distribution among 18S, 28S and other RNA classes. Benzonase-derived reads are mostly assigned to 28S rRNA, which is located at the ribosome large subunit (LSU). In contrast, RNAse I-derived reads are equally assigned to both 18S and 28S rRNAs, located at the ribosome small (SSU) and large subunit (LSU), respectively. (D) The most abundant rRNA-derived fragments obtained with Benzonase (red) and RNAse I (blue) are shown in the ribosome structure. See also Supplementary Videos 1 and 2.

**Figure 3:**
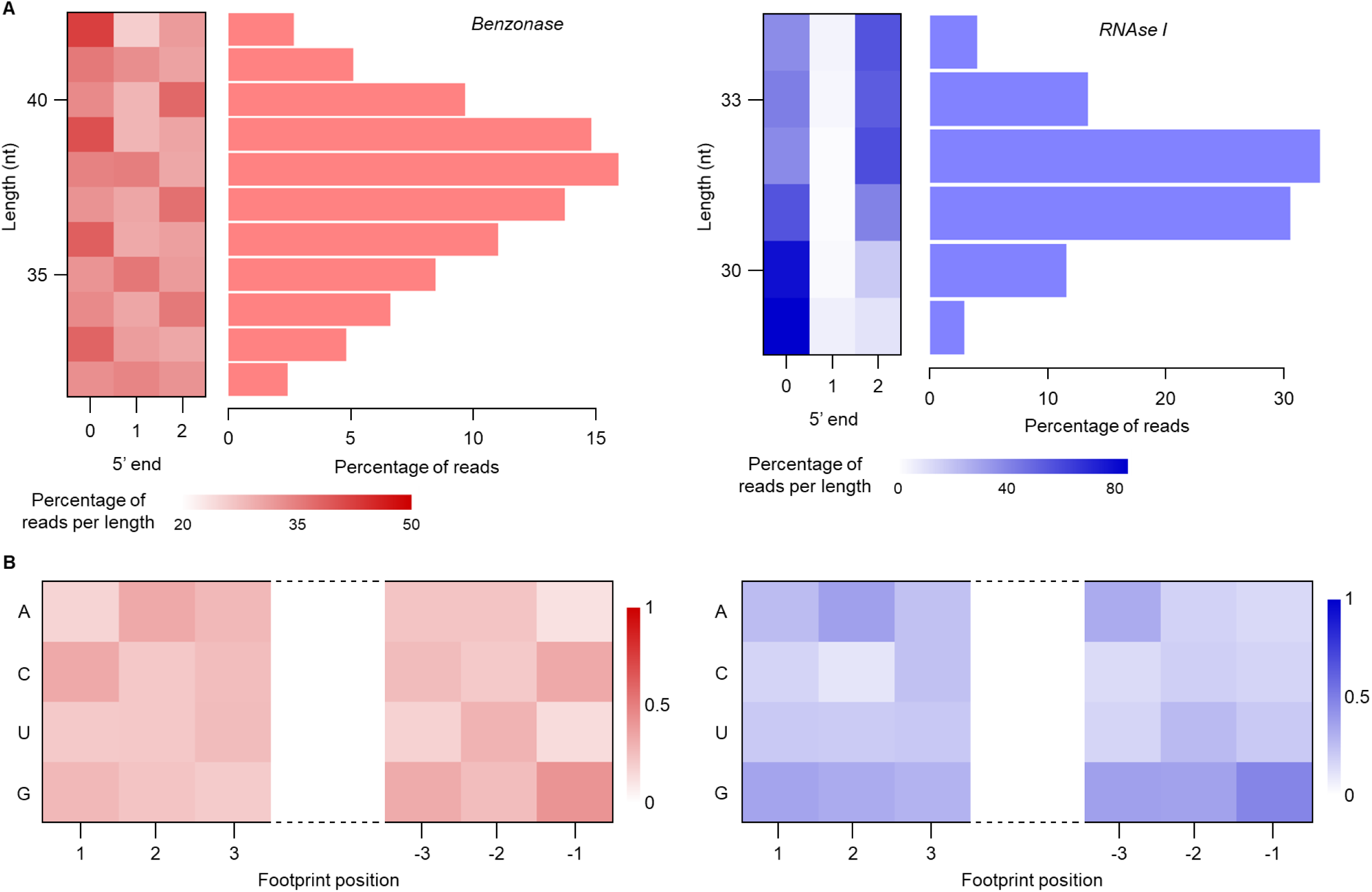
Ribosome footprints periodicity and enzyme cut bias. (A) The percentage of 5’-end ribosome footprints which are mapped in codon position 0, 1 or 2 is shown, separated by read size, for both enzymes. Footprints periodicity is observed for both enzymes but is sharper for RNAse I-derived reads. (B) Enzymes cut bias was analyzed for the first and last 3 nucleotides of the reads. No particular bias is observed for both enzymes, as expected.

We observed a pronounced mapping preference for both Benzonase- and RNAse I-derived ribosome footprints on coding regions compared to untranslated regions (Fig. 2A). Approximately 78% of Benzonase-derived footprints and 92% of RNAse I-derived footprints map to coding sequences (CDS). Although the percentage of reads mapping to 5’-untranslated region (UTR) is comparable for footprints derived from both enzymes, a distinct pattern emerges in the 3’-UTR. Specifically, Benzonase-derived footprints are more prevalent in the 3’-UTR than those from RNAse I. Prompted by this discrepancy, we investigated the read length distribution of ribosome footprints based on their mapping region (Fig. 2B). For each enzyme, ribosome footprints mapping to 5’-UTR and CDS exhibit a similar length distribution, with Benzonase-derived footprints typically centered around 37-39 nucleotides (nt) and RNAse I-derived footprints around 31-32 nt (Fig. 2B). However, footprints mapping to 3’-UTR demonstrate a different distribution, predominantly centered at 35 nt in length for both enzymes. These variations may be attributable to the distinct structural conformations adopted by the ribosome during its transit along the mRNA, especially following the completion of translation. Consequently, the observed disparity in the quantity of 3’-UTR mapping footprints between the two enzymes could be a result of differential enzyme activities on the ribosome, particularly at the 3’-UTR. To indirectly assess the extent of RNAse activity on the ribosome, we evaluated the proportion of 18S and 28S rRNA-derived fragments generated by both enzymes (Fig. 2C). Our findings reveal that RNAse I produces fragments from both rRNA classes, corresponding to the ribosomal large (LSU) and small (SSU) subunits. In contrast, Benzonase predominantly generates 28S rRNA-derived fragments specific to the LSU (Table S2 and Fig. S1). Additionally, we explored the position of the most abundant rRNA-derived fragments produced by each enzyme over the ribosome structure (Fig. 2D and Supplementary Videos 1 and 2). Concordantly with the previous observation, RNAse I-derived fragments are positioned over the large and small subunits, while Benzonase-derived fragments are only observed in the surface of the ribosome large subunit. As expected, there was minimum overlap between the fragments populations derived from these two enzymes.

We then conducted an analysis of ribosome footprint length distribution and mapping periodicity at a subcodon resolution. While Benzonase-derived footprints exhibit a broader size range (32-42 nucleotides) compared to those derived from RNAse I (29-34 nucleotides), both enzymes produce footprints demonstrating the anticipated periodicity (Fig. 3A). In Benzonase-derived footprints, the enriched subcodon position for each read size does not surpass 50%, a trend consistent with previous observations (23–26). Conversely, RNAse I-derived footprints display a characteristic subcodon distribution pattern, where the middle position is markedly underrepresented, and one of the other two positions is predominantly occupied (refer to (1, 27) for examples). Additionally, we investigated potential sequence biases of both enzymes by examining the prevalence of each ribonucleotide at the start and end of the sequenced footprints (illustrated in Figure 3B). As anticipated, neither enzyme exhibited a distinct preference for any specific ribonucleotide across the positions analyzed. However, it is noteworthy that RNAse I-derived footprints display a marked preference (approximately 50%) for guanidine residues at the 3’-end, corroborating earlier findings (9).

### RNAse I and Benzonase define comparative translatomes

The translatomes defined by each enzyme clustered as expected, according to a principal component analysis and shared a high percentage of genes (Fig. S2). A high Pearson correlation between gene expression values (Fig. 4A), as well as a similar distribution (Fig. 4B, left panel) between Benzonase- and RNAse I-derived translatomes was observed. To explore the differences between the translatomes defined by each enzyme we established a criteria to identify genes particularly overexpressed in one translatome (see Materials and Methods). Using this approach, we identified 600 mRNAs highly detected in the Benzonase-derived translatome (Benzonase-specific) and 749 mRNAs highly detected in the RNAse I-derived translatome (RNAse I-specific; Fig. 4B). In order to understand the features of these groups of mRNAs we analyzed their expression levels and the CDS length. We observed that the 600 Benzonase-specific mRNAs have a broad CDS length distribution similar to a random group (Fig. 4C, upper panel). However, the group of 749 RNAse I-specific mRNAs have a short CDS (median = 615 nt), which is even shorter than that of the first quintile of most expressed genes (usually the shorter; median = 993 nt) although the median expression of this group lies between the second and third quintile. Additionally, the size distribution of this group of 749 mRNAs differs significantly from a random group (Fig. 4C, lower panel). This suggests a potential relationship between expression level estimation and CDS length. We found that, when grouped by size, short CDS have a higher gene expression estimation in RNAse I-derived data compared to Benzonase (Fig. 4D). Conversely, large CDS appears to be more highly expressed in Benzonase-derived translatome than RNAse I-derived one, albeit to a lesser degree. As a way to extend this observation, we compared two translatome data sets derived from mouse brains obtained using either Benzonase (25) or RNAse I (28) to produce ribosome footprints. Again, we observed that short CDS have a higher expression level when using RNAse I to produce the ribosome footprints compared to Benzonase (Fig. 4D). Accordingly, the correlation between gene expression values is affected when the CDS length is incorporated into the analysis: shorter CDS showed a lower correlation, which increased with the lengthening of CDS (Fig. S3).

**Figure 4:**
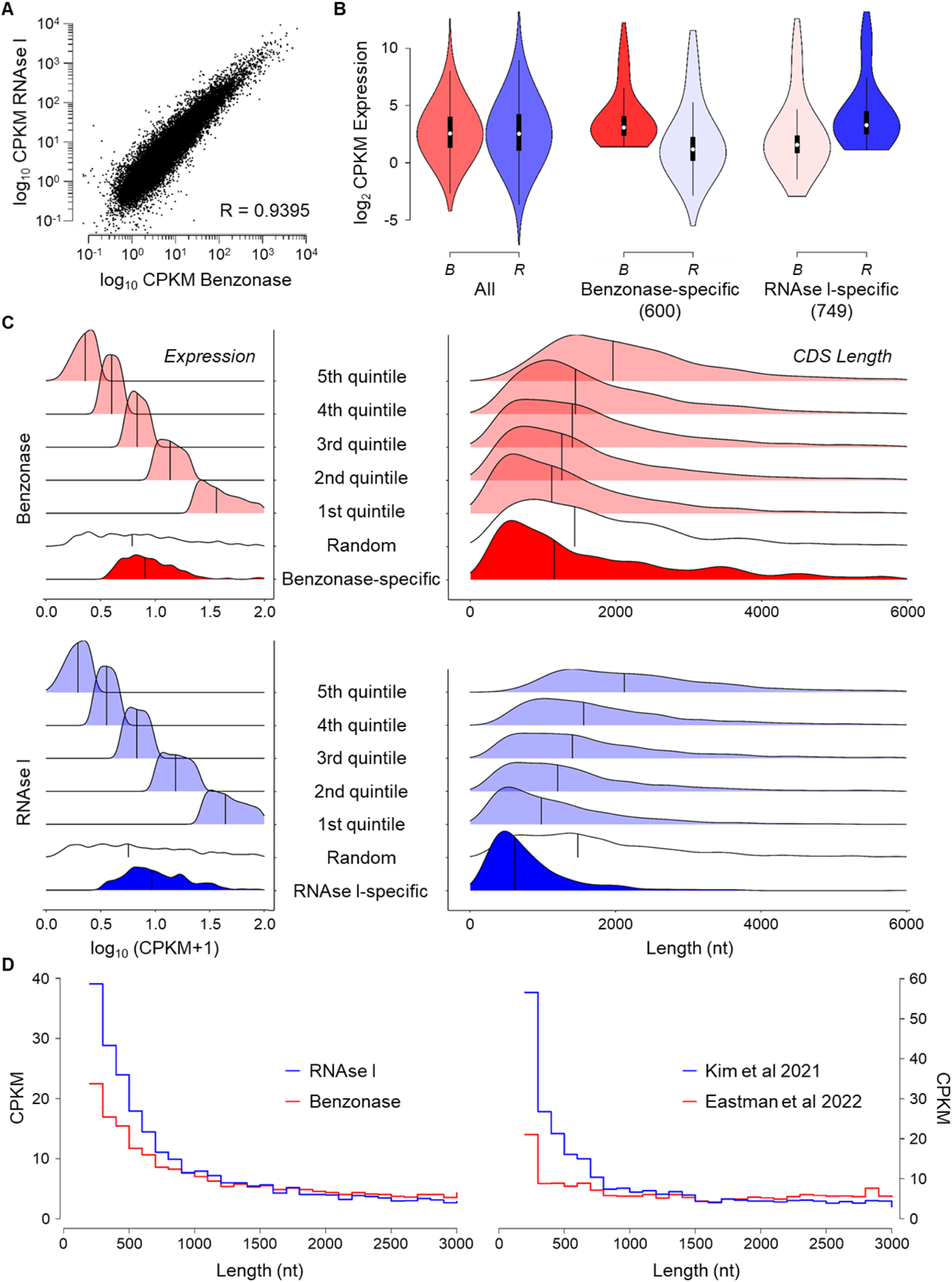
Selection and length analysis of the genes specifically detected with one enzyme. (A) Scatter plot showing translatome gene expression correlation between Benzonase- and RNAse I-derived translatomes. Pearson correlation is shown. (B) Gene expression distribution illustrated by violin plots for all mRNAs (left panel), Benzonase-specific mRNAs (middle panel) and RNAse I-specific mRNAs (right panel). (C) Expression (left) and CDS length (right) histogram distribution are shown for all the analyzed genes grouped by expression quintiles, the group of mRNAs specifically detected with one enzyme, and a size-matched random group (vertical line indicates the median). The upper panel in red indicates Benzonase-derived data and the lower panel in blue RNAse I-derived data.. (D) Median gene expression of mRNAs grouped by size (100 nt windows). Left panel shows Benzonase (red) and RNAse I (blue) translatomes. Right panel shows translatome quantification obtained from mouse brains from public data of Eastman et al. 2022 (use Benzonase, red) and Kim et al. 2021 (use RNAse I, blue).

As another approach to studying Benzonase- and RNAse I-derived translatomes, we compared the ribosome footprints mapping profiles of several highly expressed genes (Table S3), as has been done previously by others (9, 12). As expected, mapping profiles were substantially different in both translatomes, with only a handful of peaks shared (Fig. 5).

**Figure 5:**
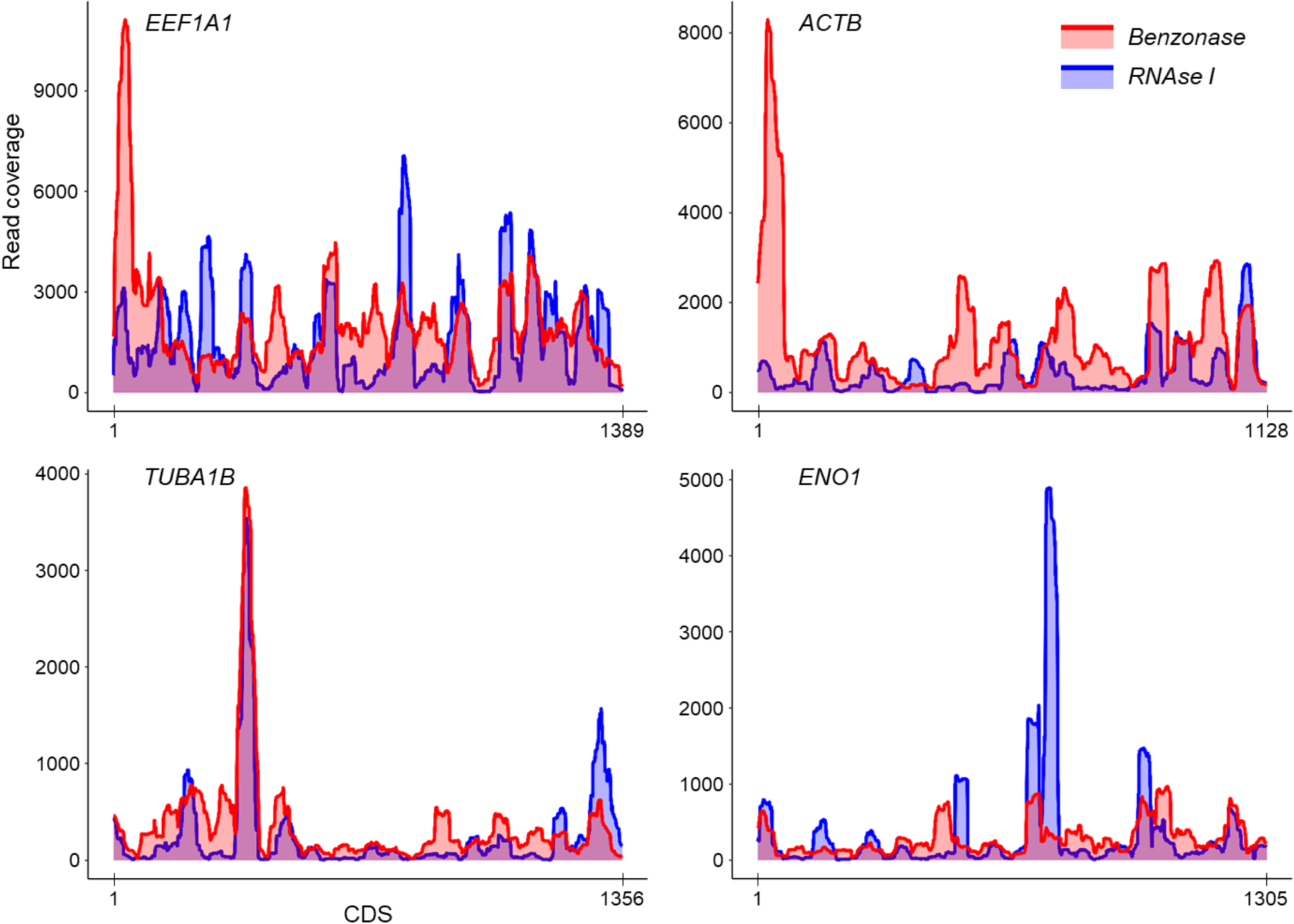
Ribosome footprint mapping coverage over highly expressed mRNAs. The mapping coverage is shown for four different translatome highly expressed mRNAs (Table S3). Benzonase-derived and RNAse I-derived mapping coverages are shown together in red and blue, respectively, for each mRNA CDS.

### Optimization of ribosome footprints production from mouse primary neuronal cultures

As an interesting model to work with and acknowledging the limitations in the amount of cells obtained, we opted to optimize the protocol of ribosome footprints production using in vitro primary neuronal cultures from embryonic mice. In this endeavor, we followed previous suggestions (7, 8) regarding the omission of the ribosome recovery step and proceeded directly with the digestion of the post-mitochondrial supernatant to subsequently recover the ribosome footprints (see Introduction). We first digested post-mitochondrial supernatant obtained from in vitro cultures of mouse embryonic neurons with different amounts of Benzonase (Fig. 6). Through this process, we were able to identify that 25 U of Benzonase is the optimal quantity to digest a post-mitochondrial supernatant derived from a 10 cm dish of mouse embryonic neurons with 4 to 6 million cells. The digestion conducted in these conditions yielded a distinct peak of about 28 nt, as expected for the ribosome footprints (Fig. 6C).

**Figure 6:**
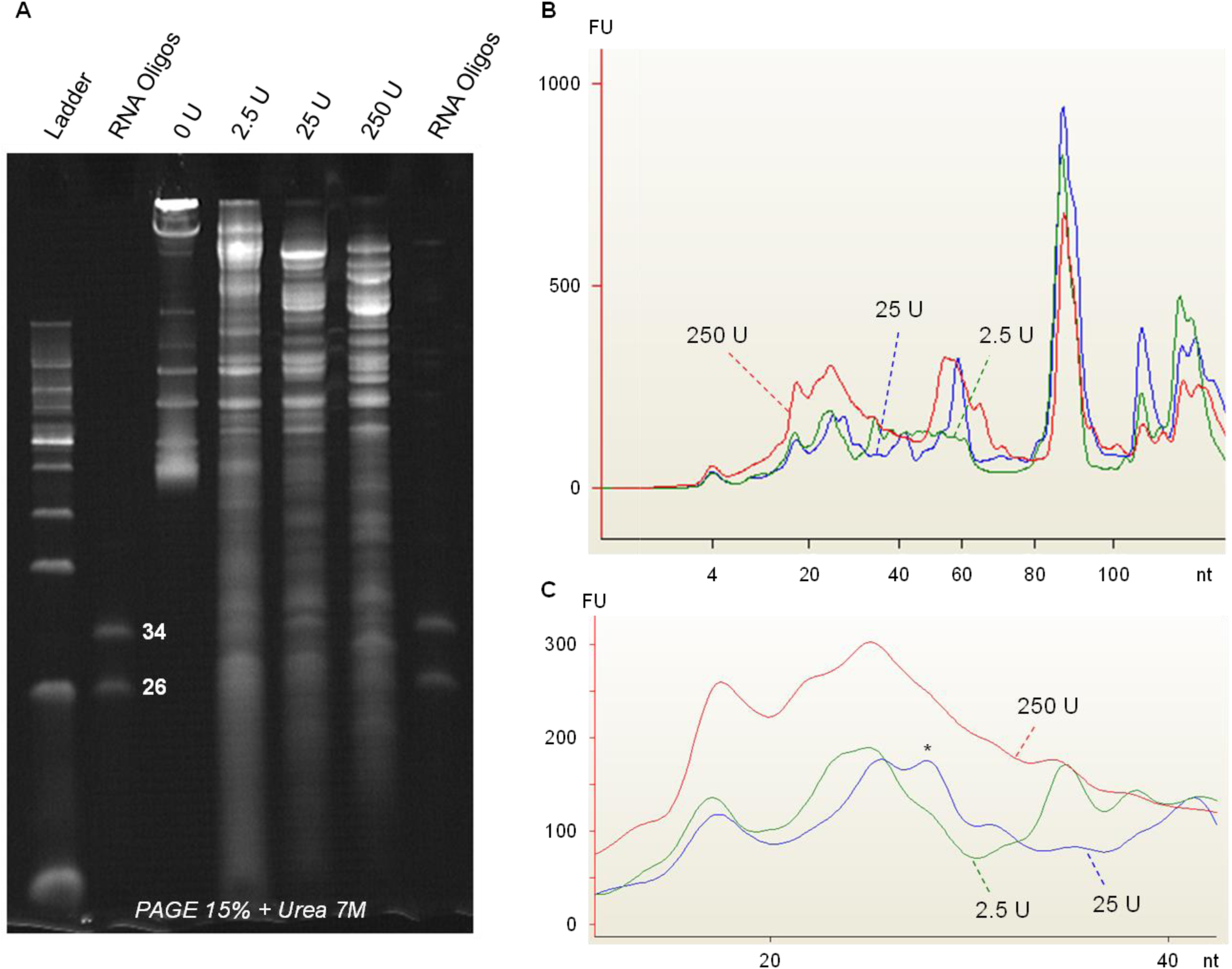
Optimization of ribosome footprints production from mouse embryonic neuronal cultures. (A) Post-mitochondrial supernatants obtained from mouse neuronal cultures were digested with different Benzonase amounts (0 -no digestion-; 2.5; 25 and 250 units) and separated in 15% PAGE + 7M Urea. ssRNA oligos of 34 and 26 nt in length were used to indicate the ribosome footprints region. (B) Samples were separated using capillary electrophoresis (Agilent - Bioanalyzer small RNA chips) and the electropherogram profile is shown for each digestion in order to visualize RNA population. (C) The electropherogram ribosome footprints region (20-40 nt) is zoom-in and shown for the three digestions. An asterisk (*) indicates the 28 nt peak that corresponds to the ribosome footprints’ standard size.

We then applied this protocol to produce ribosome footprints from mouse neuronal cultures derived from wild-type (WT) and tau knock-out (tauKO) mice (Fig. S4). On average, we obtained 116 million reads, of which 81% corresponded to rRNA. Non-rRNA reads were mapped to the reference mouse transcriptome, yielding about 11 million mapped reads. We analyzed several global features of the resulting translatomes to assess their quality (Fig. 7). Initially, we evaluated the number of detected genes across increasing cutoffs. As expected, we detected almost 10, 000 genes with at least 10 CPM (Fig. 7A). The global distribution of gene expression levels appears as anticipated, and no differences were observed between the two used genotypes (Fig. 7B). We also investigated the distribution of ribosome footprints across mRNAs features (Fig. 7C). As observed earlier (Fig. 2A), the ribosome footprints are mainly located in CDS regions, but some of them are found in the 3’-UTR. Similarly, footprint length distribution and read periodicity were studied in the two data sets derived from both genotypes. Again, as expected for ribosome footprints obtained using Benzonase (23–26), the length of the reads range between 30 to 40 nt and periodicity levels were close to 50% (Fig. 7D and E).

**Figure 7:**
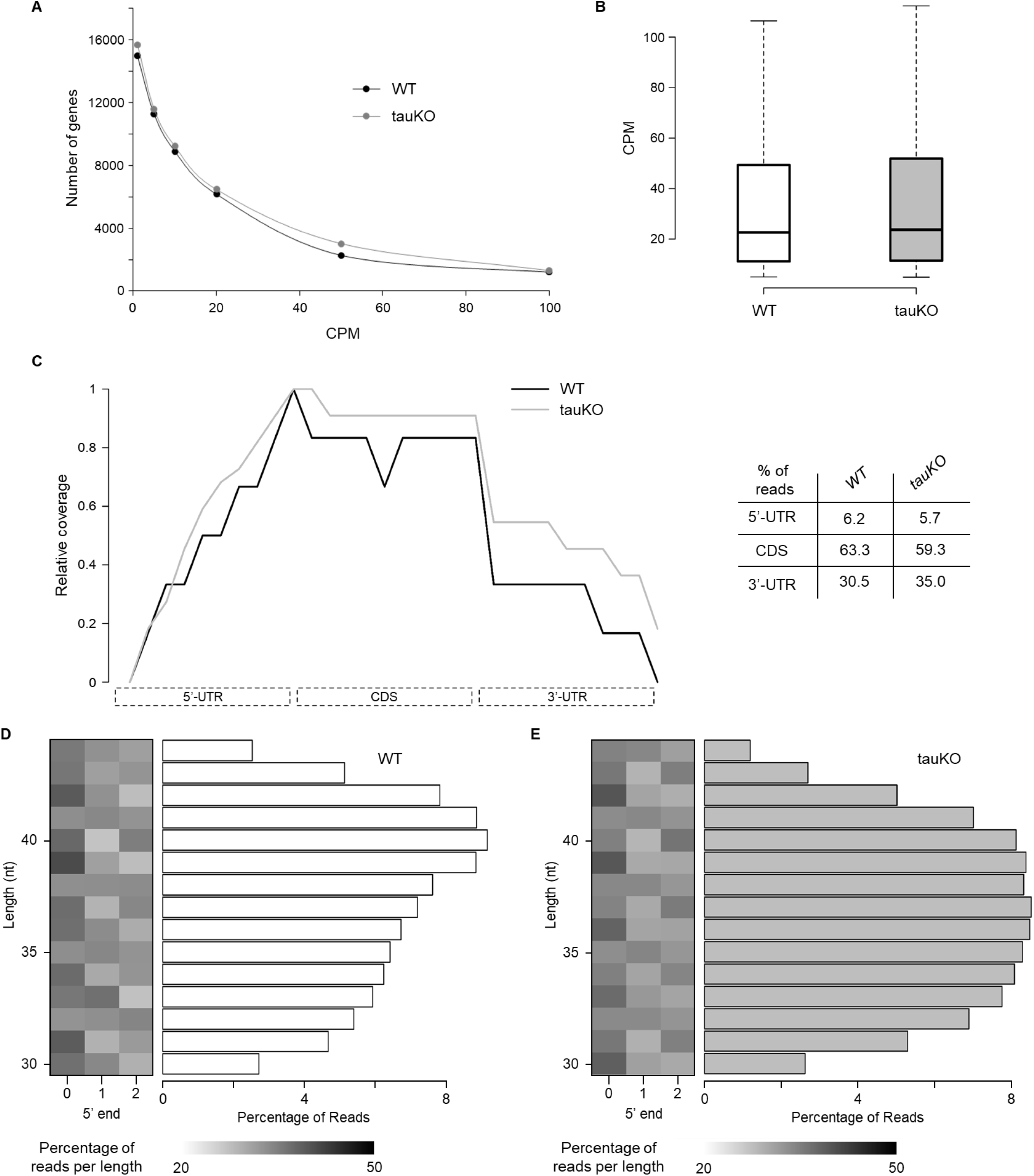
Main features of the mouse neuronal cultures derived translatomes. Ribosome footprints were obtained by post-mitochondrial supernatant digestion from primary neuronal cultures derived from wild-type (WT) and tau knock-out (tauKO) mice. (A) The number of observed genes at different CPM cutoffs is shown for both genotypes. About 10, 000 genes show an expression level of at least 10 CPM. (B) Global gene expression distribution is shown for both genotypes as boxplots. (C) Coverage plot of ribosome footprints over mRNA regions for both genotypes. The percentage of reads mapping in each region is also shown as a table. For both genotypes, footprints are particularly abundant in coding sequences. (D and E) As shown in Figure 3, the percentage of 5’-end ribosome footprints which maps in codon position 0, 1 or 2 is shown, separated by read size, for both genotypes. Footprint periodicity is observed in both cases.

## DISCUSSION

Ribosome profiling, or Ribo-seq, is a genomic tool for the study of translation at a genome-wide level with sub-codon resolution. Since its publication in 2009, this technique has been widely expanded with only minor modifications in the original protocol, mainly associated with the library preparation steps. However, the RNAse protection assay has been raised by some groups as a critical step in ribosome footprint production (9–11). In this work, our objective was to investigate the use of Benzonase, an enzyme that has not been explored previously, to produce ribosome footprints. Until now, the RNAse I enzyme has been the most commonly employed, as described in the original protocol (4). This non-specific nuclease of 27 kDa was first isolated from E. coli and it is specific for RNA, particularly single-strand tracts (29, 30). However, a few other enzymes have been described and used in the literature (9, 12). In our study, we optimized and utilized the Benzonase nuclease (23–26, 31).

This nuclease, also known as Serratia endonuclease, is obtained from Serratia marcescens and consists of two subunits of 30 kDa (32, 33). Like RNAse I, Benzonase is non-specific but can degrade RNA and DNA as single or double strands. Importantly, as mentioned in the Introduction, Benzonase leaves the 5’ and 3’ fragment ends ready for ligation steps, minimizing protocol complexity, reducing bias sources and increasing yield.

In order to compare both enzymes, we obtained polysome samples derived from HEK293 cell cultures and digested them, in parallel, using Benzonase and RNAse I enzymes. The ribosome footprints obtained with both enzymes were subjected to high-throughput sequencing and we compared several footprints’ features as well as the quantified translatomes for each enzyme. As expected, each enzyme produces a unique digestion pattern with a specific fragment population in terms of quantity and size (Fig. 1). We then explored features such as mapping distribution over mRNAs, length and periodicity (Fig. 2 and 3). Despite the enzyme used, the mRNA coding sequence is the region where it is most probable to find a mapping footprint. However, we found about 16% of Benzonase-derived footprints mapping in the 3’-UTR (Fig. 2A), as was also previously observed for micrococcal nuclease (MNase)-derived footprints (12). Interestingly, when we compared the length of the footprints in relation to the mRNA region where they had mapped, we found that 3’-UTR-mapped footprints exhibited a distinct length distribution compared to 5’-UTR or CDS footprints (Fig. 2B), as was also previously reported for MNase-derived footprints (12). Based on the hypothesis that the presence of ribosome footprints on 3’-UTR could be due to differences in enzyme activity, we indirectly evaluated the activity of both enzymes on the ribosome by exploring the proportion of 18S and 28S rRNA-derived fragments produced by each enzyme. We found that RNAse I produces rRNA fragments derived from both 18S and 28S classes, while Benzonase only produces 28S-derived rRNA fragments (Fig. 2C). This could indicate a strong RNAse I activity over the ribosome, resulting in the production of rRNA fragments derived from both ribosome subunits, whereas Benzonase exhibits a more gentle activity that exclusively produces rRNA fragments derived from the large subunit (Fig. 2D). The hypothesis of differential enzyme activity over the ribosome aligns with the observed results in read periodicity (Fig. 3). In this context, Benzonase would produce footprints with a broader length distribution and a more diffuse pattern of periodicity. In contrast, a tight degradation by the RNAse I would yield fragments of specific sizes with a clear periodicity, as was observed here and discussed by others (11).

We also investigated differences and commonalities between the translatomes obtained from the quantification of ribosome footprint counts over the transcriptome using either Benzonase or RNAse I. As expected, we only found minor global differences between both translatomes (Fig. 4A and 4B, Fig. S2). However, individual mRNAs exhibit distinct mapping patterns (Fig. 5), as previously documented (9). We then applied filter criteria to detect those genes specifically identified, or highly detected, with one enzyme but not the other (see Material and Methods). The most noteworthy characteristic of the groups of specific genes was the length of their CDS (Fig. 4C). RNAse I-specific genes have significantly shorter CDS lengths compared to the overall CDS length distribution, while conversely, Benzonase-specific genes display a broader length distribution. We then explored if this bias was also present at a genome-wide level and found that genes with short CDS seem to be particularly highly detected in RNAse I data. This phenomenon was also observed in previously published data sets derived from a different organism (Fig. 4D). This observation can be explained, as mentioned before, in terms of the differential enzyme activity hypothesis over the ribosome. Short CDS, usually highly expressed, would have a high ribosome density that can be resolved by a small enzyme, such as RNAse I, that will produce individual ribosome footprints. On the other hand, the Benzonase enzyme (about two times the size of RNAse I) will face more issues digesting ribosome-crowded mRNAs. The same argument will explain why long CDS are usually less expressed in RNAse I-derived data compared to Benzonase-derived: RNAse I would be able to over-digest long CDS with fewer ribosomes, while Benzonase would produce representative footprints.

In the last place, following the published protocol alternatives (7, 8) we adopted our Ribo-seq protocol to generate the ribosome footprints directly from the post-mitochondrial supernatant, thereby avoiding the most sample-demanding step of ultracentrifugation (Fig. 6). By using this alternative approach, we were able to produce and sequence ribosome footprints derived from mouse embryonic neuronal cultures (Fig. S4). As previously analyzed, the obtained ribosome footprints and translatomes exhibit the expected features for the two genotypes utilized in this study (Fig. 7). This example expands the range of biological models for which the Ribo-seq technique can be applicable, addressing a particular challenge in the context of neuronal-related models.

In sum, this work represents a detailed comparison between the ribosome footprints produced with Benzonase and RNAse I, and between the translatomes defined by each set of footprints. The significance of the enzyme selection for Ribo-seq has been highlighted previously by others (9), and a small group of enzymes has been studied and characterized, especially the MNase (12). However, this work represents, to the best of our knowledge, the first detailed characterization of ribosome footprints obtained with Benzonase. The comparison with RNAse I-derived footprints revealed interesting results and clear differences in terms of length, periodicity pattern and distribution among mRNA’s features, which we hypothesize could be explained by disparities between enzyme activities over the RNA. We acknowledge certain caveats associated with the use of Benzonase, such as the generation of larger footprints with a broader length distribution and a weaker periodicity pattern compared to RNAse I-derived footprints. These differences may limit some downstream analyses, including P-site occupancy, codon bias, and translational pause detection. Nevertheless, we believe that Benzonase offers a significant advantage by simplifying the protocol, reducing processing time, and minimizing potential biases relative to RNAse I, making it a valuable alternative enzyme for translatome quantification by Ribo-seq. Finally, we observed that several features of the Benzonase-derived footprints have been also recognized in MNase-derived footprints (12). In this line, alternative enzymes capable of producing ribosome footprints will be powerful tools for the community as the Ribo-seq technique becomes more popular and different protocols emerge. One clear example is the recent description of a single-cell version of the Ribo-seq protocol that uses MNase to produce the ribosome footprints (34), highlighting the need to expand our knowledge of different and new enzymes.

## ACKNOWLEDGEMENTS

We would like to thank Dr. Nutan Shivange for handling mice and preparing primary neuron cultures, and members of the Sotelo-Silveira lab for the discussing the results and providing intellectual input. We also want to sincerely thank Dr. David Munroe (†), former director of the Laboratory of Molecular Technologies at LEIDOS, Frederick National Laboratory for Cancer Research, NCI. His original contributions and fundamental ideas during the early stages of this project laid the foundation for this work.

## AUTHOR CONTRIBUTIONS

Guillermo Eastman: Conceptualization, Data curation, Formal analysis, Investigation, Methodology, Validation, Visualization, Writing—original draft. George S. Bloom: Investigation, Resources, Funding acquisition, Writing—review & editing. José R. Sotelo-Silveira: Conceptualization, Investigation, Resources, Supervision, Funding acquisition, Writing—review & editing.

## SUPPLEMENTARY DATA

Supplementary Data and Videos 1 and 2 are available at NAR online.

## CONFLICT OF INTEREST

The authors have no conflicts of interest to report.

## FUNDING

This work was supported by the PhD fellowships program from Agencia Nacional de Investigación e Innovación (ANII) [POS NAC 2016 1 129959 to G.E.]; PROLAB travel grant (PABMB/ASBMB/IUBMB) and The Pew Charitable Trusts through a Pew Latin American Fellows Program in the Biomedical Sciences fellowship to G.E; Programa de Desarrollo de las Ciencias Basicas (PEDECIBA) to G.E. and J.R.S-S.; National Institutes of Health [AG051085 to G.S.B.]; Owens Family Foundation; Cure Alzheimer’s Fund; and the Rick Sharp Alzheimer’s Foundation to G.S.B.

## DATA AVAILABILITY

The data underlying this article are available in the NCBI Sequence Read Archive (SRA) at https://www.ncbi.nlm.nih.gov/sra, and can be accessed with BioProject ID PRJNA1220030 and PRJNA1220074.

## Notes

### Competing Interest Statement

The authors have declared no competing interest.

