## Supplementary Data for "The use of Benzonase to produce ribosome footprints simplifies translational levels quantification by Ribo-seq"

### MANUSCRIPT TITLE

### SUPPLEMENTARY DATA

This document includes Supplemental Figures 1 to 4 and Supplemental Tables 1 to 3.

**SUPPLEMENTARY FIGURES**

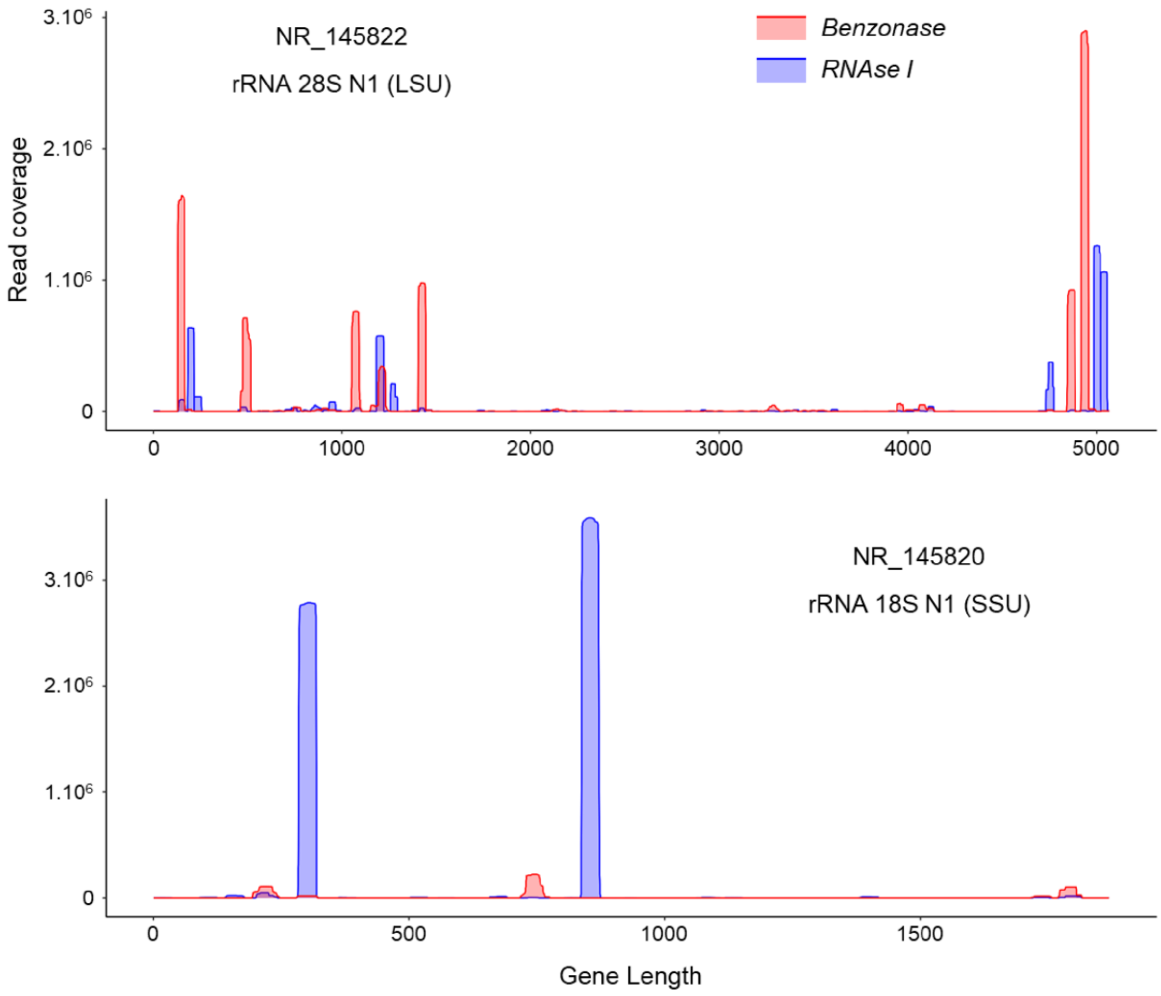


**Figure S1: Mapping profile of reads over rRNAs.** Mapping profiles of reads derived from Benzonase (red) and RNAse I (blue) treatments are shown for the rRNA 28S N1 gene (NR_145822, upper panel) and the rRNA 18S N1 gene (NR_145820, lower panel).

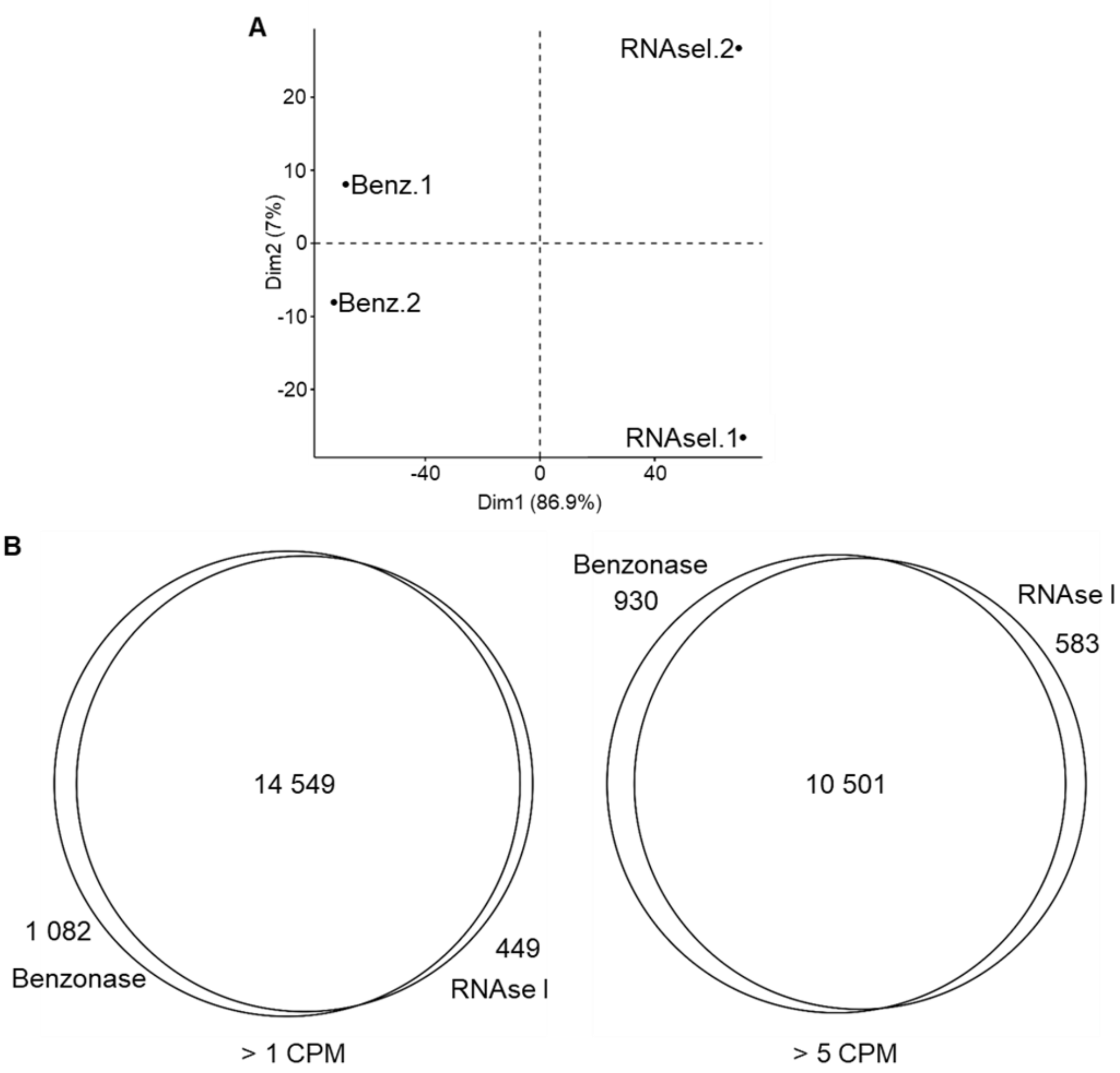


**Figure S2: Global comparisons between Benzonase- and RNAse I-derived translatomes.** (A) Principal component analysis of all translatome samples. (B) Venn diagrams showing the number of shared genes in Benzonase- and RNAse I-derived translatomes at different cutoffs of >1 CPM (left) and >5 CPM (right).

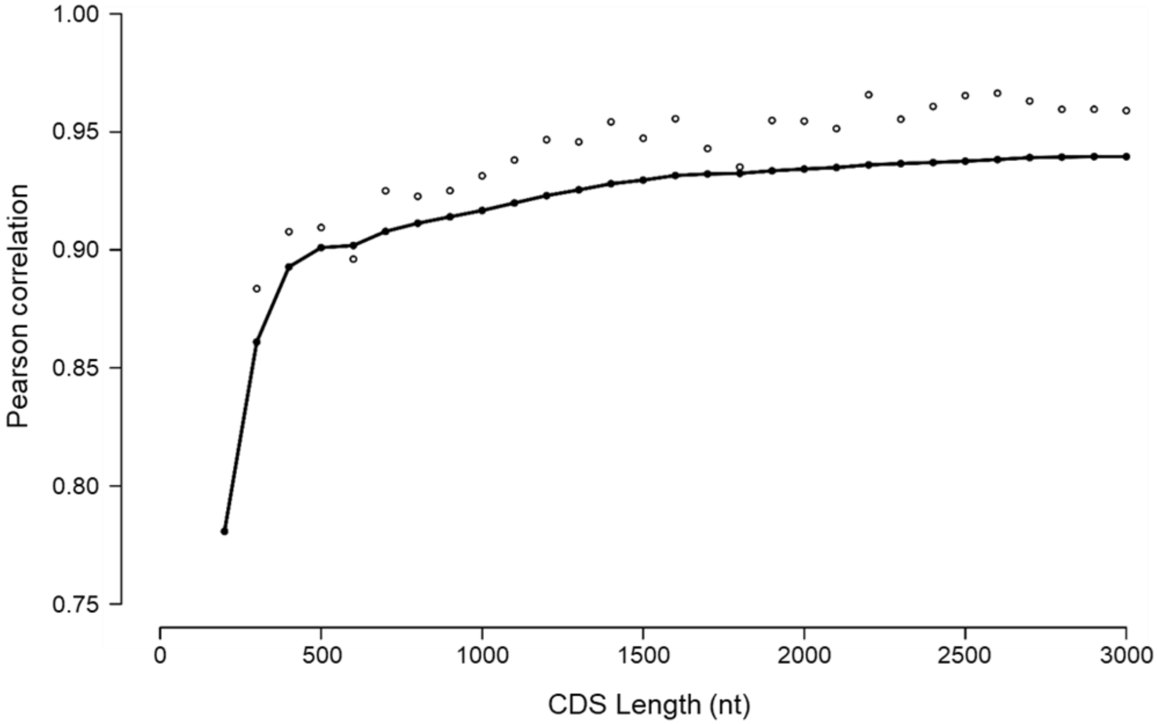


**Figure S3: Pearson correlation between Benzonase- and RNAse I-derived translatomes grouping mRNAs by size.** Individual dots show Pearson correlation value for each subgroup of mRNAs ordered by size using a 100 nt window from 200 to 3000 nt. The line shows the accumulative Pearson correlation value.

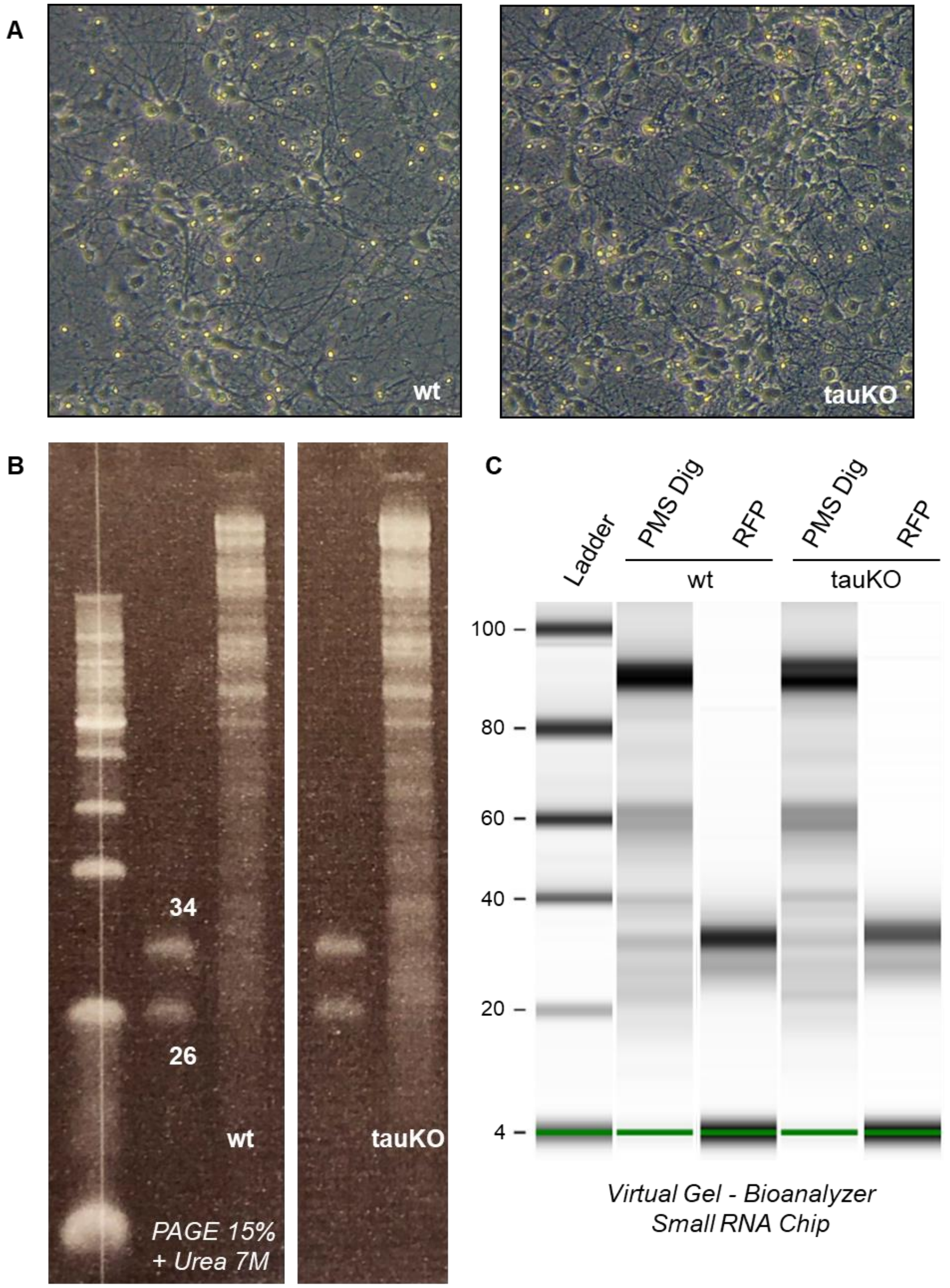


**Figure S4: Ribosome footprints production from in vitro neuronal primary cultures.** (A) Illustrative images of the neuronal cultures after 10-11 DIV for both of the genotypes used (WT - wild type; and tauKO - tau knock-out). (B) Ribosome footprints separation and isolation from 15% PAGE + 7M urea. From left to right: Molecular weight DNA ladder, ssRNA oligos, WT neurons digestion, ssRNA oligos, tauKO neurons digestion. (C) A virtual reconstruction of the gel, obtained from the Agilent Bioanalyzer capillary electrophoresis, is shown for both post-mitochondrial supernatant digestions (PMS Dig) and isolated ribosome footprint samples (RFP) derived from both genotypes (WT and tauKO).

### SUPPLEMENTARY TABLES

**Table S1:** The number of raw sequenced reads is shown for each sample along with rRNA-derived reads, genome- and mRNA-mapped reads.

| Enzyme | rep # | Raw Reads | rRNA | | Genome mapped reads | mRNA mapped reads | |
|---|---|---|---|---|---|---|---|
| Benzonase | 1 | 122,906,986 | 100,574,709 | 81.8% | 16,673,224 | 14,720,802 | 88.3% |
| | 2 | 95,768,276 | 80,593,765 | 84.2% | 9,438,694 | 8,106,113 | 85.9% |
| RNAse I | 1 | 123,579,208 | 108,402,139 | 87.7% | 8,549,450 | 7,310,863 | 85.5% |
| | 2 | 153,015,842 | 135,748,360 | 88.7% | 10,700,276 | 9,222,608 | 86.2% |

**Table S2:** Sequence and size of the observed rRNA-derived fragments produced by each enzyme.

| Benzonase | |
|---|---|
| 28 rRNA-derived fragments | Size |
| 5'-CCGCGGCGGGGCGCGGGACATGTGGCGTACGGAAGACC-3' | 38 |
| 5'-GGATTCAACCCGGCGGCGGGTCCGGCCGTGTCGGCGGCCCGGCGGATCTTTCCC-3' | 54 |
| 5'-CGGCGTCTCCTCGTGGGGGGGCCGGGCCACCCCTCCCACGGCGCGAC-3' | 47 |
| 5'-CGCGCTCGCCGGCCGAGGTGGGATCCCGAGGCCTCTCCAG-3' | 40 |
| 5'-CGCGCGCCGGGACCGGGGTCCGGTGCGGAGTGCCCTTCGTCCTG-3' | 44 |
| 5'-CCCTCGCCCGTCACGCACCGCACGTTCGTGGGGAACCTGG-3' | 40 |
| RNAse I | |
| 28 rRNA-derived fragments | |
| 5'-TCGTGGGGGGCCCAAGTCCTTCTGATCGAGGCC-3' | 33 |
| 5'-CAGTGCGCCCCGGGCGGGTCGCGCCGTCGGGCCCGGGGG-3' | 39 |
| 5'-AGCGCCGCGGAGCCTCGGTTGGCCTCGGATAGCCGGTCCCCCG-3' | 43 |
| 5'-CTGGGTCGGGGTTTCGTACGTAGCAGAGCAGCTCC-3' | 35 |
| 5'-TCGCTGCGATCTATTGAAAGTCAGCCCTCGACACAA-3' | 36 |
| 18 rRNA-derived fragments | |
| 5'-CTTTGGTGACTCTAGATAACCTCGGGCCGATCGCAC-3' | 36 |
| 5'-GCCGCCTGGATACCGCAGCTAGGAATAATGGAATAG-3' | 36 |

**Table S3:** The number of reads and ranking position is shown for the five highly expressed genes selected for shown ribosome footprints mapping coverage patterns.

| Gene | ID | CDS Length | Benzonase | | RNAse I | |
|---|---|---|---|---|---|---|
| | | | Ranking | Reads | Ranking | Reads |
| EEF1A1 | NM_001402 | 1,389 | 1 | 77,973 | 2 | 71,624 |
| ACTB | NM_001101 | 1,128 | 6 | 35,930 | 39 | 15,842 |
| ENO1 | NM_001428 | 1,305 | 82 | 10,343 | 60 | 15,006 |
| TUBA1B | NM_006082 | 1,356 | 52 | 13,251 | 105 | 10,821 |
